# BSAseq: an interactive and integrated web-based workflow for identification of causal mutations in bulked F2 populations

**DOI:** 10.1101/2020.04.08.029801

**Authors:** Liya Wang, Zhenyuan Lu, Michael Regulski, Yinping Jiao, Junping Chen, Doreen Ware, Zhanguo Xin

## Abstract

**Summary:** With the advance of next-generation sequencing (NGS) technologies and reductions in the costs of these techniques, bulked segregant analysis (BSA) has become not only a powerful tool for mapping quantitative trait loci (QTL) but also a useful way to identify causal gene mutations underlying phenotypes of interest. However, due to the presence of background mutations and errors in sequencing, genotyping, and reference assembly, it is often difficult to distinguish true causal mutations from background mutations. In this study, we developed the BSAseq workflow, which includes an automated bioinformatics analysis pipeline with a probabilistic model for estimating the segregation region and an interactive Shiny web application for visualizing the results. We deeply sequenced a male sterile parental line (*ms8*) to capture the majority of background mutations in our bulked F2 data. We applied the workflow to 11 bulked F2 populations and identified the true causal mutation in each population. The workflow is intuitive and straightforward, facilitating its adoption by users without bioinformatics analysis skills. We anticipate that BSAseq will be broadly applicable to the identification of causal mutations for many phenotypes of interest.

**Availability:** BSAseq is freely available on https://www.sciapps.org/page/bsa

## 1 Introduction

Bulked segregant analysis (BSA) is a QTL mapping technique for identifying genetic markers associated with a mutant phenotype (Michelmore *et al*., 1991). For BSA, individuals are usually selected from the tails of the phenotypic distribution from a genetic cross to form two pools (bulks) of segregants. Then, both pools are genotyped to identify the genomic region containing the causal loci, for which allele frequency should differ between the two bulks.

Ethyl methanesulfonate (EMS) mutagenesis has been widely used to introduce novel mutations into plants such as *Arabidopsis* to reveal the functions of many genes (Page and Grossniklaus, 2002). Coupling EMS mutagenesis with the NGS technique, MutMap (Abe *et al*., 2012) has been proposed as a means for identifying causal loci in rice by backcrossing an EMS mutant with a trait of interest to the wild-type parent and then sequencing the segregant F2 pools. The concept of MutMap is theoretically straightforward. Because the mutant phenotype is selected from an F2 segregation population, the allele frequency of the causal mutation should be 100%. In other words, the SNP ratio (the number of short reads with the mutation divided by the total short reads covering the site) should be 1. By contrast, the SNP ratio of mutations unlinked to the causal mutation should be around 0.5, and the SNP ratio of linked mutations should vary from around 0.5 to 1 depending on the genetic distance to the causal mutation.

For functional validation of genes in sorghum, we generated 6400 pedigreed M4 mutant pools (with 253 pools sequenced) from EMS-mutagenized BTx623 seeds (Jiao *et al*., 2016). By coupling BSA with NGS, we identified the functions of several recessive sorghum genes using bulked F2 pools derived by backcrossing EMS mutants with a wild-type BTx623 founder line (BTx623′). These genes include *msd*1 (Jiao, Lee, *et al*., 2018), *bm40* (Jiao, Burow, *et al*., 2018), *msd*3 (Dampanaboina *et al*., 2019), and *ms9* (J. Chen *et al*., 2019). To filter out false-positive candidates, the causal genes were usually identified using more than one EMS mutant. Furthermore, for a mutation to be considered, an allele frequency of 1.0 was strictly required; this criterion made the detection of causal mutations sensitive to insufficient sequencing depth and errors in genome assembly, alignment of short reads, genotyping, or phenotyping.

In this study, we improved and integrated a BSAseq workflow into a cloud-based workflow platform, SciApps.org (Wang *et al*., 2018), which can be used by thousands of users through NSF-funded resources including the CyVerse Data Store (Goff *et al*., 2011) for storage, XSEDE resources (Towns *et al*., 2014) at the Texas Advanced Computing Center (TACC) for computing, and a federation system (Wang, *et al*., 2015) at the Cold Spring Harbor Laboratory (CSHL) for data management, visualization, and additional computing. The workflow supports the rapid identification of causal genes with an automated bioinformatics analysis pipeline, a probabilistic model for quantitatively scoring segregation regions, and a Shiny web application for interactively visualizing analysis results directly on the SciApps platform. To validate the improved workflow, we demonstrated that all causal genes discovered in the studies cited above could be correctly identified with BSAseq using one or at most two F2 pools.

## 2 Methods

### 2.1 Overview

Fig. 1 (A) depicts the process of using BSAseq to discover the causal genes for a trait of interest. We begin by backcrossing the mutant that has the desired phenotype with the wild type BTx623′. The F1 plants should have the wild-type phenotype, and in F2 plants the segregation ratio between mutant and wild-type phenotypes should be approximately 1:3. The mutant-type F2 plants are pooled together for sequencing. Short reads are aligned to the reference BTx623 assembly using Bowtie2 (Langmead and Salzberg, 2012), and SNPs are called and filtered using Bcftools (Li, 2011) to keep only EMS-induced mutations (G → A or C → T) with the desired coverage (e.g., 5–100). SnpEff (Cingolani *et al*., 2012) is used to annotate and select mutations with a large predicted effect (missense, nonsense, splice_site_acceptor, and splice_site_donor), and SIFT 4G (Vaser *et al*., 2016) is used to predict whether these mutations are deleterious or not. Each of the above analytical steps is integrated into SciApps as a modular application (app) and chained together as an automated pipeline; an automatically generated diagram is shown in Fig. 1 (B). The first two apps are configured to run on the TACC resources, the other apps are executed on the CSHL federation system, and all results are archived in the CyVerse Data Store.

**Fig. 1.**
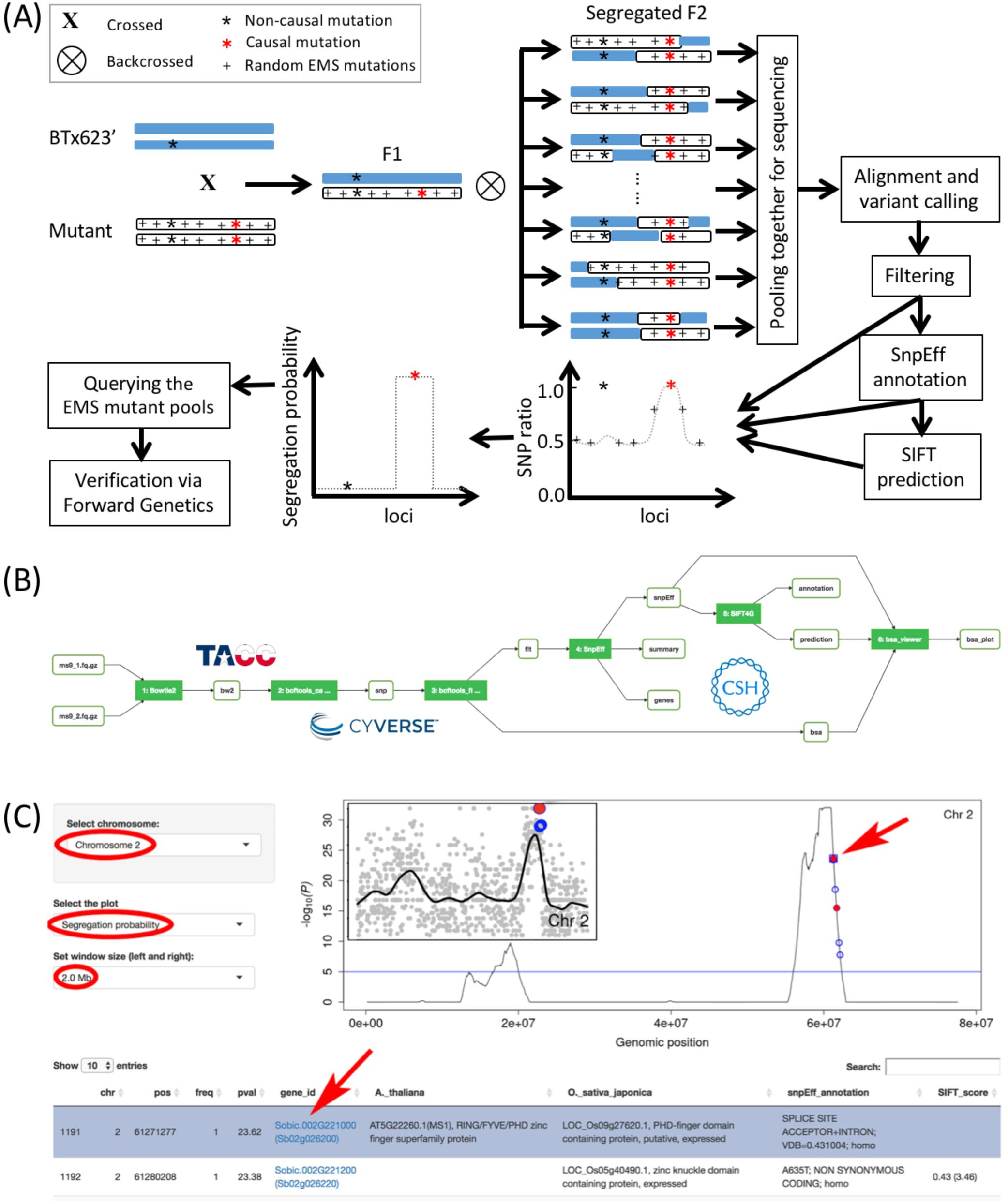
The BSAseq workflow. (A) Overview. (B) Diagram of the analysis pipeline chaining files (open button) and applications (filled buttons). (C) Segregation probability plots with a window size of 2 Mb along the chromosome 2 for the ms9 F2 pool data. Blue circles indicate nonsynonymous SNPs. Filled red circles indicate significant mutations (stop_gained, splice_site_acceptor, splice_site_donor, start_lost, or missense mutations with SIFT score < 0.05 and median_info < 3.25). The blue horizontal line indicates the 10^−5^ significance threshold. The first row of the table (below the plot) is selected by clicking, which highlights the row in blue and the matching mutation on the plot with a blue square (pointed by a red arrow). Note that clicking on any circles on the plot will also highlight the matching genes in the table. Clicking on the candidate gene id will query the EMS mutation database and return the mutation line with independent significant mutation(s) in the same gene. The inset figure is the SNP ratio plot of chromosome 2 where each gray dot represents a SNP and the solid line is the result of LOWESS (LOcally WEighted Scatter-plot Smoother) regression.

As shown by the diagram, the final app of the pipeline, bsa_viewer, aggregates the results from the previous three steps to support interactive visualization. When users attempt to visualize the output of the app, the results will be copied from the CyVerse Data Store to the CSHL federation system (if they are not already stored there), and an interactive Shiny web application will be opened. As a demonstration, we sequenced an F2 pool from a cross between a male-sterile mutant isolated from the pedigreed line Mut574 and the wild type BTx623′. The causal gene (the red dot with the lowest *P*-value) falls in the segregated region on chromosome 2 (Fig. S1). The data can also be viewed by the SNP ratio. Most importantly, biological knowledge can be also applied to eliminate false-positive candidate genes (or genes with the candidate mutations). The detailed calculations of the segregation probability are described in the next section. The raw sequence data of Mut574 have been deposited into NCBI SRA (PRJNA580273).

To run the workflow in sorghum or another organism, only a few modifications are needed: 1. Select the paired sequencing data through “Browse” in step 1; 2. Select the required reference genome in steps 1, 2, 4, and 5; 3. Specify your own or select the background SNPs in step 3. Default settings can be used for all other parameters. A detailed tutorial is available: https://tinyurl.com/rgc5tjz. Data should be uploaded to the CyVerse Data Store via CyberDuck (https://cyberduck.io/) or iCommands (https://docs.irods.org/4.2.1/icommands/user/). A free CyVerse account is required to host sequencing data at CyVerse. After a user runs BSAseq, SciApps will archive the analysis results in their CyVerse account and chain the raw data and analysis results together as shareable automated workflows.

### 2.2 Estimating the segregation probability

As shown in Fig. 1 (A), the causal mutation for the trait of interest is expected to have an SNP ratio of 1 (or close to 1 if there are any errors in the data collection or from analysis tools). For mutations close to the causal loci, the SNP ratio should decrease from 1 with increasing genetic distance, whereas unlinked mutations are expected to have ratios around 0.5. However, the SNP ratio plot can be noisy due to unfiltered background mutations, as shown in Fig. S2 and the inset figure of Fig. 1 (C). To improve the signal-to-noise ratio, we used a weighted one-sample single-tailed t-test to estimate the significance of an EMS mutation falling within the segregated region.

The null hypothesis is that the weighted mean of the SNP ratios around the mutation (+/-*W*) is equal to 0.5 (no segregation); *W* is the window size. The weighted mean and standard deviation are defined by equations (1) and (2), with the weight *w*_*i*_ defined by equation (3), a tricube kernel function. *D*_*i*_ is the standardized distance, with value 0 at the focal position and value 1 at the edge of the window. *N* is the number of EMS mutations in the window, *M* is the number of nonzero weights, and *x*i is the SNP ratio.

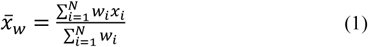

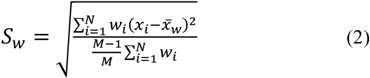

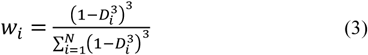

The logarithm of the *P*-values is plotted along each chromosome by the Shiny web application, as shown in Fig. S1, to easily visualize the segregated region as well as the true-positive candidate genes, which should fall into the segregated region.

### 2.3 The Shiny web application

The weights defined by the tricube kernel function (equation (3)) are more critical when a larger window size is needed, e.g., when the segregated region is broad due to a low rate of recombination. We created a Shiny web application to support interactive the window size; toggling of the view between SNP ratio and segregation probability; aggregation of homologous gene annotation (*A. thaliana* and *O. sativa japonica*), SnpEff annotations, and SIFT score predictions; and querying of previously identified EMS mutations (Jiao *et al*., 2016) (Fig. 1 (C)).

After completing the BSAseq analysis pipeline, users can open the Shiny web application by visualizing the bsa_viewer output. In the beginning, segregation probability plots along all chromosomes are displayed (Fig. S1). Users can choose different window sizes to improve the estimation of the segregated region. Once a chromosome with a segregated region is determined, users can zoom into the chromosome to more closely examine the candidate genes (Fig. 1 (C)). SNPs are automatically annotated in the workflow, allowing users to determine the causal mutation based on biological knowledge. The interactive Shiny web application can toggle from the statistic graph to the traditional SNP ratio graph for confirmation of the causal SNP, which should have a ratio of nearly 1 and should lie within a region devoid of unlinked genetic mutations (e.g., the peak on chromosome 2 in the inset figure of Fig. 1(C)).

Once a candidate gene is identified, the user can click the gene id to get the EMS mutant that has a significant mutation on the same gene by querying the EMS mutation database. The database is built with the previously sequenced 253 EMS mutant pools (Jiao *et al*., 2016) to help identify the independent allele of the causal mutation for confirmation of predicted candidate genes. We are in the process of sequencing an additional 1000 mutant pools to cover the majority of sorghum genes.

### 2.4 Deep sequencing of the sorghum *ms8* mutant line for capturing the background mutations

As shown in Fig. S2, many SNPs far away from the causal loci have a SNP ratio near 1, e.g., the two false-positive SNPs (red dots) on chromosome 5. The reasons why these SNPs have ratios close to 1 include low sequencing coverage around the mutations, local segregation distortion, and most importantly, differences between the reference BTx623 and the founder BTx623′ line used to create the EMS mutants; these mutations will have SNP ratios of 1 because they are present in both parents. In Fig. 1 (A), a heterozygous SNP in the founder line is used as an example to show that false-positive candidate genes can be generated by insufficient sequencing or sampling.

Additionally, sorghum is primarily self-pollinated, meaning that an individual sorghum plant will accept pollen from its own flowers. To ensure that we are backcrossing the EMS mutant with BTx623′, we used the *ms8* mutant (Xin *et al*., 2017), which was a male sterile mutant isolated from the sorghum mutant library and has been backcrossed to the parent BTx623′ for six generations, as the wild-type parent for creating F2 BSA mapping populations. Because *ms8* is completely male-sterile, all seeds derived from the crosses of mutants to *ms8* are truly F1 seeds. We deeply sequenced *ms8* to 60X coverage and discovered 12,675 EMS-type (GC ? AT) mutations. After subtracting these background mutations, candidate mutations were located in one chromosome region, greatly simplifying the calling of the causal mutations. The raw sequencing data of *ms8* were deposited into NCBI SRA (PRJNA606537).

## 3 Results

A male-sterile mutant isolated from the mutant line Mut574 was mapped to chromosome 2 based on the statistical model (Fig. S1), The peak contained two mutations predicted to be deleterious (red dots) and three mutations with tolerable effects (blue circles). Flipping of the graph to the SNP ratio (Fig. S2) revealed that these mutations are located on the tip of a region devoid of unlinked genetic mutations. Examination of the SNP effect of the mutations revealed that one deleterious mutation was a splice-site mutation in the gene *Sobic*.*002G221000*, a homolog of *Arabidopsis* male sterile 1 (*ms1*). This mutant happened to be allelic to the recently published *ms9* gene (J. Chen *et al*., 2019).

We applied the workflow to 11 published MutMap populations identified through conventional BSA analysis of NGS sequencing data of bulked F2 populations. This led to the identification of the correct causal mutations in each case, as shown in Fig. S3 (*ms9* (J. Chen *et al*., 2019)), S4 (*msd1* (Jiao, Lee, *et al*., 2018)), S5 (*bm40* (Jiao, Burow, *et al*., 2018)), and S6 (*msd3* (Dampanaboina *et al*., 2019)). The results are summarized in Table 1.

**Table 1.** Benchmark results for published sorghum genes

| Mutant | Causal gene | #FP<sub>*ms8*</sub><sup>a</sup> | #FP<sub>BSA</sub><sup>b</sup> | #Candidates (with TP)<sup>c</sup> | TP detected w/o filtering *ms8*<sup>d</sup> |
| --- | --- | --- | --- | --- | --- |
| Mut574 | *ms9* | 1 | 3 | 2 (Y) | Y |
| BC1F2 | *ms9* | 1 | 3 | 3 (Y) | Y |
| p9 | *msd1* | 0 | 0 | 2 (Y) | Y |
| p12 | *msd1* | 1 | 1 | 3 (Y) | Y |
| bm40-1 | *bm40* | 0 | 2 | 2 (Y) | Y |
| bm40-2 | *bm40* | 2 | 1 | 4 (Y) | Y |
| p6A | *msd3* | 1 | 1 | 2 (Y) | Y |
| p6B | *msd3* | 1 | 1 | 2 (Y) | N |
| p14 | *msd3* | 0 | 0 | 2 (N)<sup>e</sup> | N |
| p21A | *msd3* | 2 | 2 | 3 (Y) | Y |
| p21B | *msd3* | 1 | 2 | 3 (Y) | N |
| p24 | *msd3* | 1 | 2 | 3 (Y) | Y |

Table 1 shows that BSAseq can filter out about 50% of the false-positive candidate genes, but more than one candidate gene was left for each F2 mutant pool due to linkage to the causal gene. Fortunately, the EMS mutation sites are mostly randomly distributed, so the causal gene can be identified using another EMS mutant with an independent significant allele in the same gene. As an example, the only candidate gene shared by Mut574 and BC1F2 of *ms9* is *Sobic*.*002G221000*. Our results also showed that *msd1, bm40*, and *msd3* could all be identified with at most two F2 pools. The confounding issue for identification of *msd3* was a background mutation from *ms8* that is present in five of the six F2 pools.

BSAseq failed to detect the causal gene for p14 due to a slightly larger median info value predicted by SIFT4G, although the causal mutation did fall into the predicted segregation region. Therefore, users need to pay attention to false-negative candidate mutations if they are in the segregation region but just fail to meet the SIFT criteria for being considered as significant mutations. The *ms8* parental line not only introduces false-positive candidate genes (though all of them are outside the predicted segregation region for the BSA data listed in Table 1, and can therefore be filtered out by BSAseq) but also decreases the signal-to-noise ratio, enabling BSAseq failed to predict the segregation region for p6B and p21B (plots not shown).

Interestingly, when the p9 data were aligned to v3, the SNP ratio for the causal gene, *msd1*, was 0.96 (vs. 1 when aligned to v2). This suggests that the causal gene could not be detected if we continue to exclude SNPs with a SNP ratio less than 1.

## 4 Discussion

BSA analysis of bulked NGS data often requires a comprehensive understanding of bioinformatic tools, impeding the wide application of this technology for causal gene identification. By contrast, the simplicity of our workflow allows users who lack training in bioinformatics to run analyses. With this workflow, it is possible to identify the causal mutation with 15x coverage of the genome. In sorghum (genome size ∼700 Mb), 10 Gb of clean data, which now costs less than $200 including library construction, is sufficient to identify the causal mutations as long as the phenotype is accurate. Additionally, the workflow was developed to identify causal mutations for recessive nuclear mutations. For dominant mutations, both heterozygous and homozygous mutants yield the same mutant phenotype; consequently, the segregated region will have an average SNP ratio of 0.75. Our workflow correctly predicts the segregation region and identifies the true causal gene, e.g., for the red root mutant (results not shown; patent and publication are pending).

Genome editing technologies promise to transform breeding in both plants (K. Chen *et al*., 2019) and animals (Tait-Burkard *et al*., 2018). To effectively apply genome-editing tools, however, it is essential to know the target gene to be edited, or ideally the causal mutations. As sequencing technologies continue to be developed and sequencing costs continue to drop, BSAseq will increasingly become an affordable means to discover targets for genome editing when a high-quality mutant library is available. Induced mutant populations often have a high density of background mutations, which impedes the immediate utility of traits identified from mutant libraries in breeding. Although recurrent backcrosses can be used to remove unlinked background mutations, it will take several generations to remove 90% of them. Furthermore, no effective method currently exists for removing the linked mutations by backcrossing. On the other hand, genome editing can introduce precise mutations with few or no offsite mutations, allowing the rapid introduction of superior traits into elite germplasm for breeding. The combination of affordable and fast target gene discovery using BSAseq with precise genome editing has the potential to revolutionize breeding.

## 5 Conclusion

We demonstrated the utility of a web-based BSAseq workflow in the identification of causal mutations underlying phenotypes of interest. We showed that false-positive candidate genes can be effectively eliminated if they are outside the predicted segregation region, which can be estimated with adjustable window sizes via the interactive Shiny web application. The BSAseq workflow is species-agnostic, and could readily be applied to identifying causal genes in other plant species. We expect that the BSAseq workflow will be a useful tool for large-scale identification of causal mutations in sorghum and other organisms.

## Acknowledgments

We thank the CyVerse project, the XSEDE project at TACC, and CSHL for providing computational resources.

## Disclaimer

Mention of trade names or commercial products in this article is solely for the purpose of providing specific information and does not imply recommendation or endorsement by the U.S. Department of Agriculture. USDA is an equal opportunity provider and employer.

