## Supplementary file for "BSAseq: an interactive and integrated web-based workflow for identification of causal mutations in bulked F2 populations"

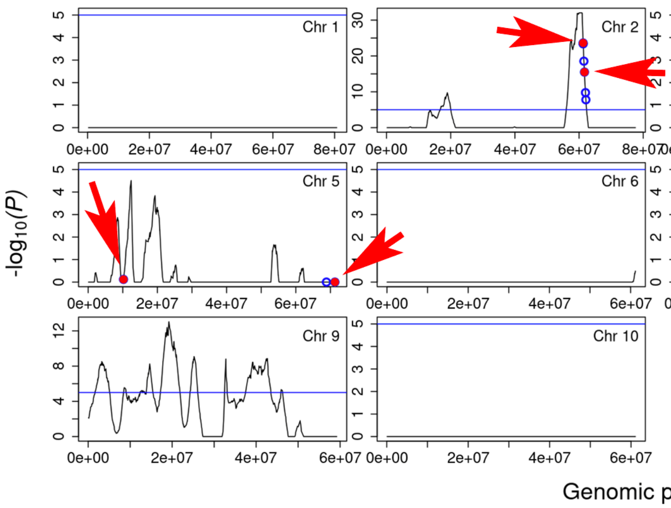


**Figure S1. Segregation probability plots with a window size of 2 Mb along the chromosomes for the *ms9* F2 pool data**.

Blue circles indicate nonsynonymous SNPs. Filled red circles indicate significant mutations, including stop_gained, splice_site_acceptor, splice_site_donor, start_lost, or missense mutations with SIFT score < 0.05 and median_info < 3.25. The blue horizontal line is each chromosome panel indicates the $10^{-5}$ significance threshold. Red arrows point to all significant mutations.

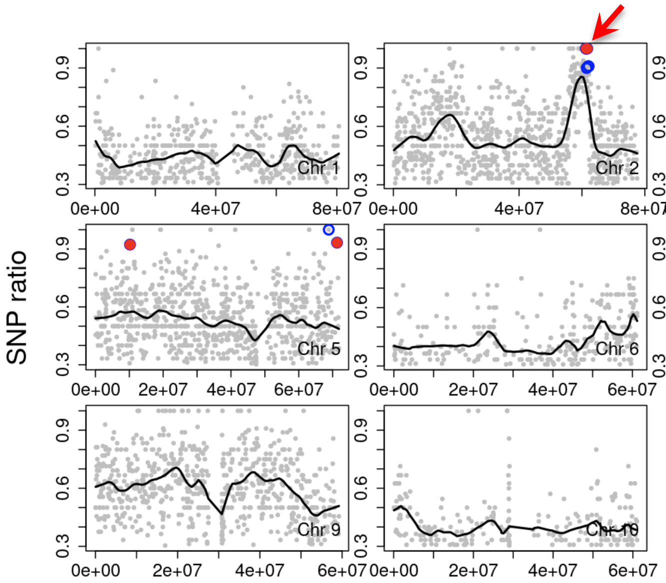


**Figure S2. SNP ratio plots along the chromosomes for the *ms9* F2 pool data**.

Each gray dot represents one SNP. Blue circles are nonsynonymous SNPs. Filled red circles indicate significant mutations, including stop_gained, splice_site_acceptor, splice_site_donor, start_lost, or missense mutations with SIFT score < 0.05 and median_info < 3.25. The red arrow points to the causal mutation. Each gray dot represents a SNP and the solid line is the result of LOWESS (LOcally WEighted Scatter-plot Smoother) regression.

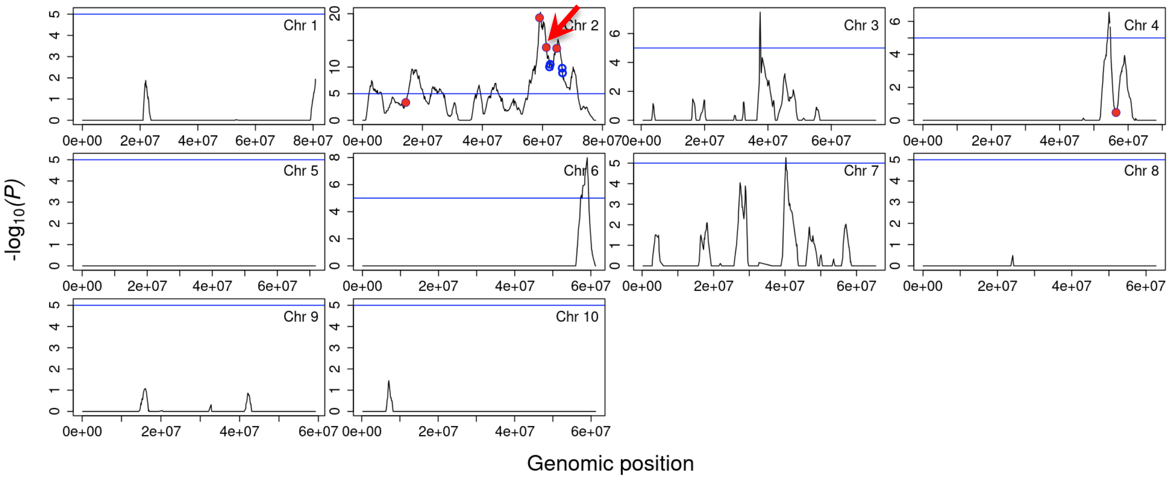


**Figure S3. Segregation probability plots with a window size of 2 Mb along the chromosomes for the *ms9* F2 pool BC1F2**.

The red arrow points to the causal mutation.

(A)

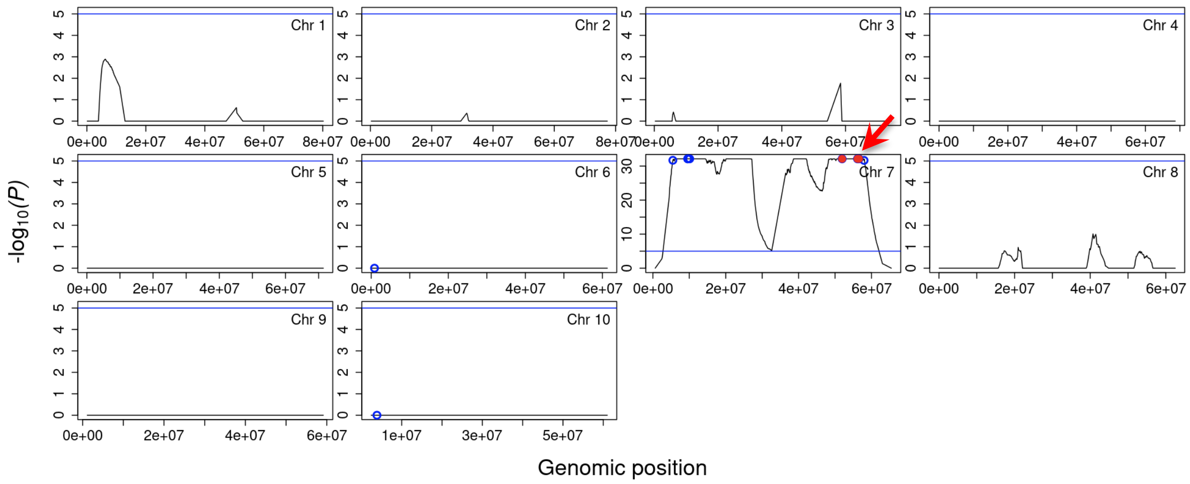


(B)

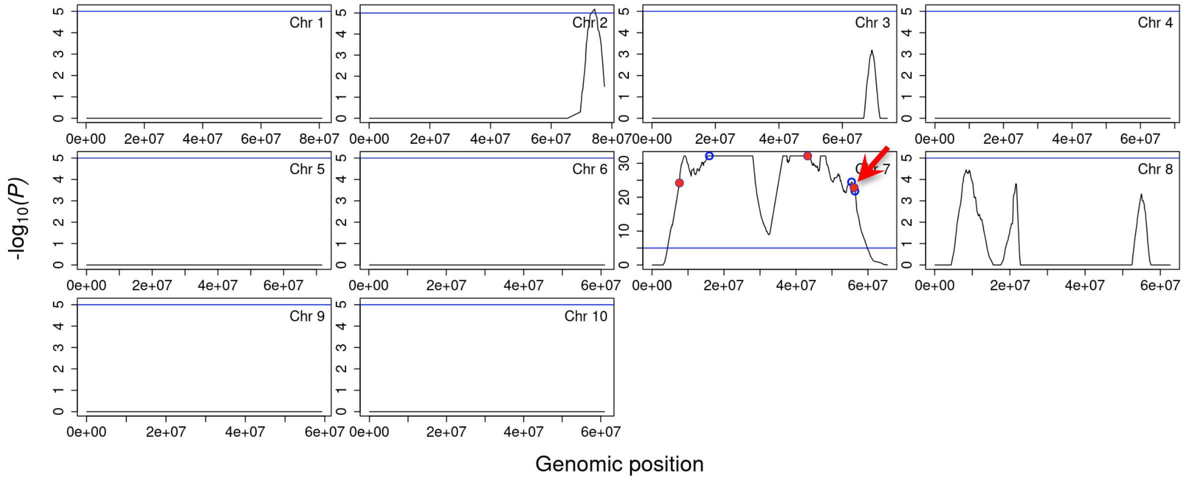


**Fig. S4 Segregation probability plots with a window size of 5 Mb along the chromosomes for the *msd1* F2 pools p9 (A), and p12 (B)**.

The red arrow points to the causal mutation.

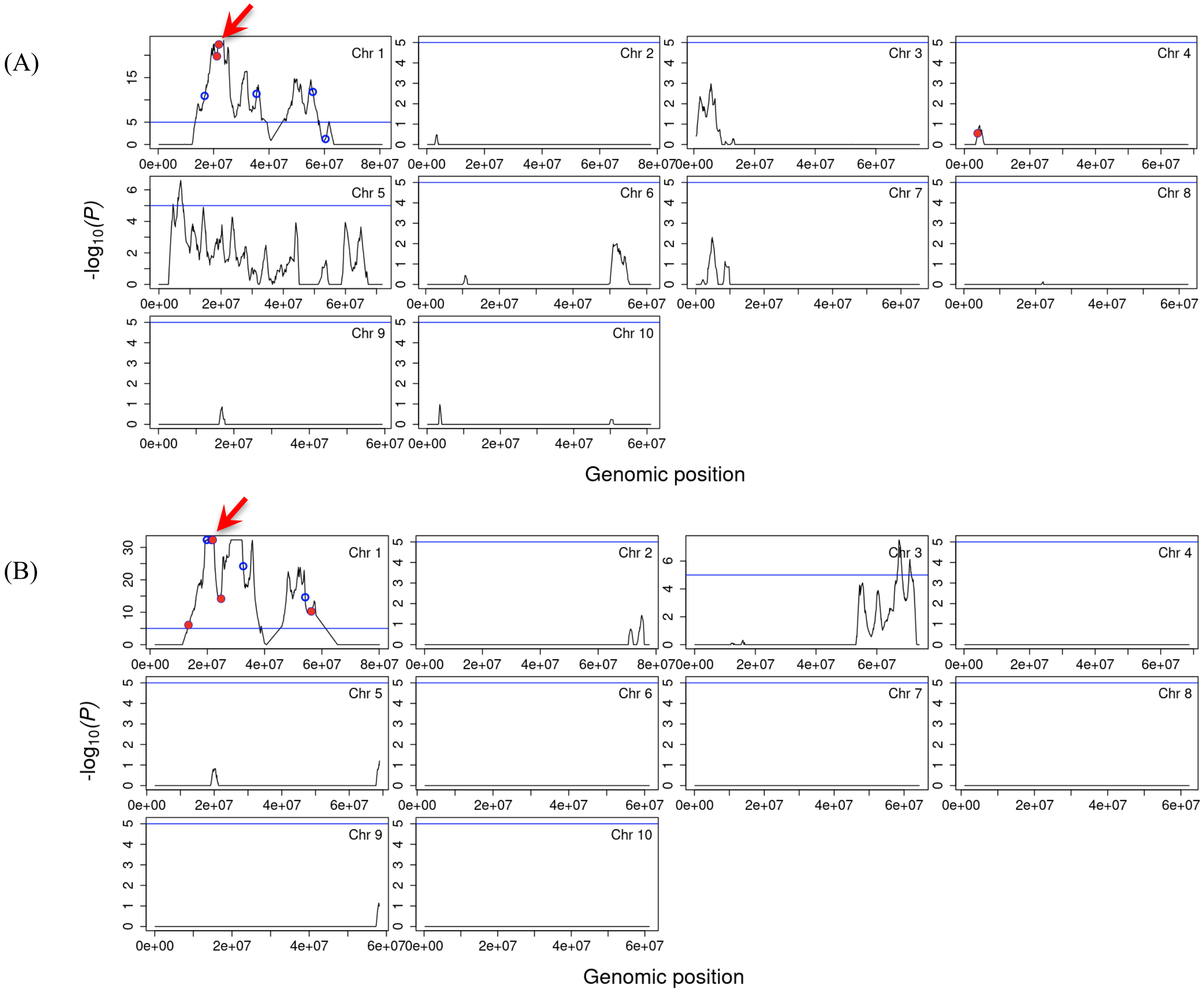


**Fig. S5 Segregation probability plots with a window size of 2 Mb along the chromosomes for the *bm40* pools bm40-1 (A), and bm40-2 (B)**.

The red arrow points to the causal mutation.

(A)

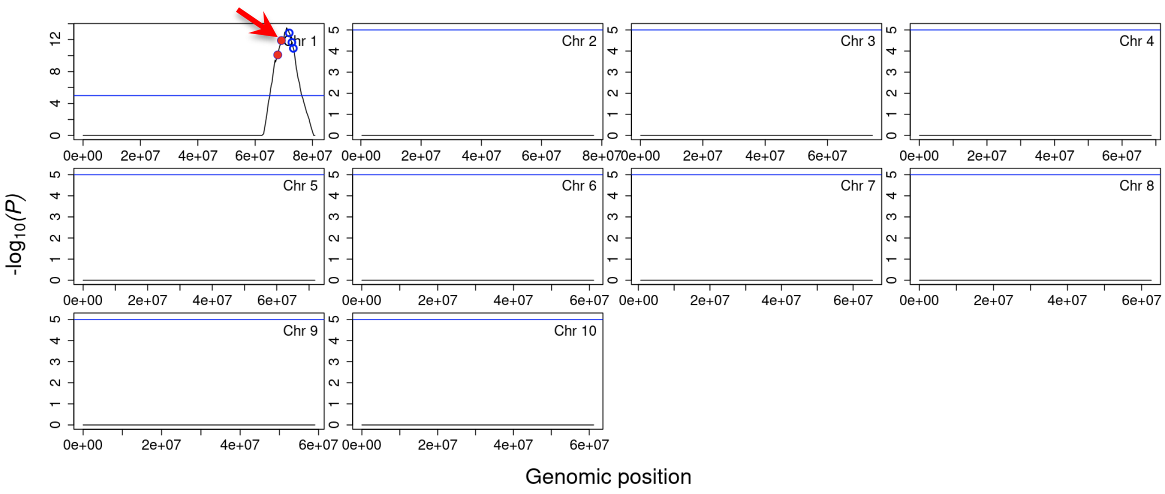
Chr 1
Chr 2
Chr 3
Chr 4
Chr 5
Chr 6
Chr 7
Chr 8
Chr 9
Chr 10
-log10(P)
Genomic position

(B)

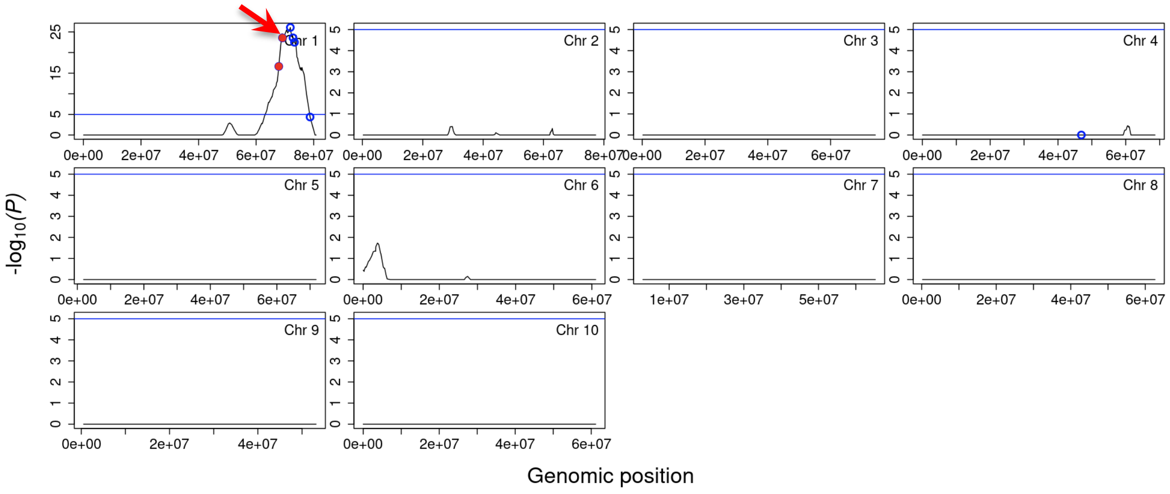
Chr 1
Chr 2
Chr 3
Chr 4
Chr 5
Chr 6
Chr 7
Chr 8
Chr 9
Chr 10
-log10(P)
Genomic position

(C)

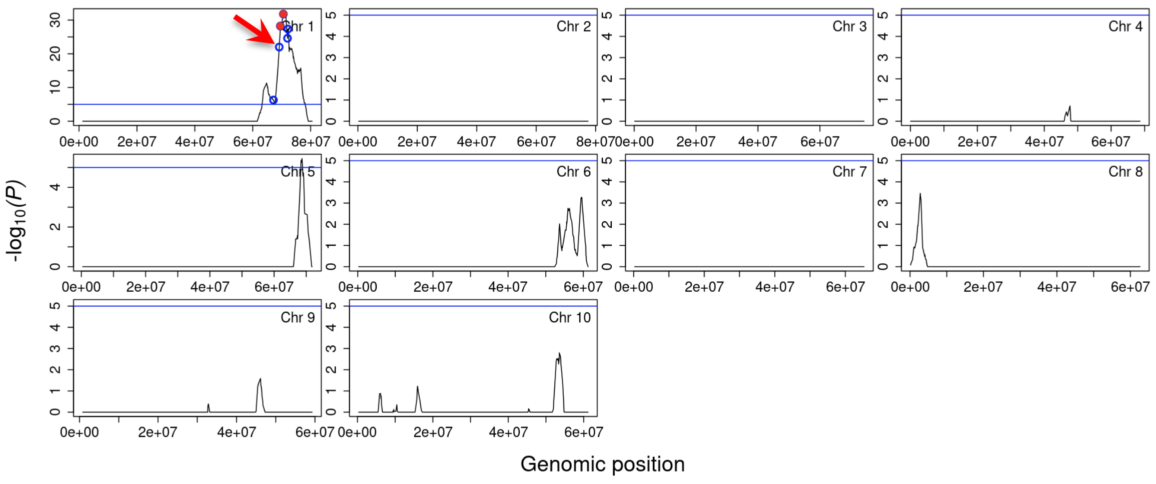
Chr 1
Chr 2
Chr 3
Chr 4
Chr 5
Chr 6
Chr 7
Chr 8
Chr 9
Chr 10
-log10(P)
Genomic position

(D)

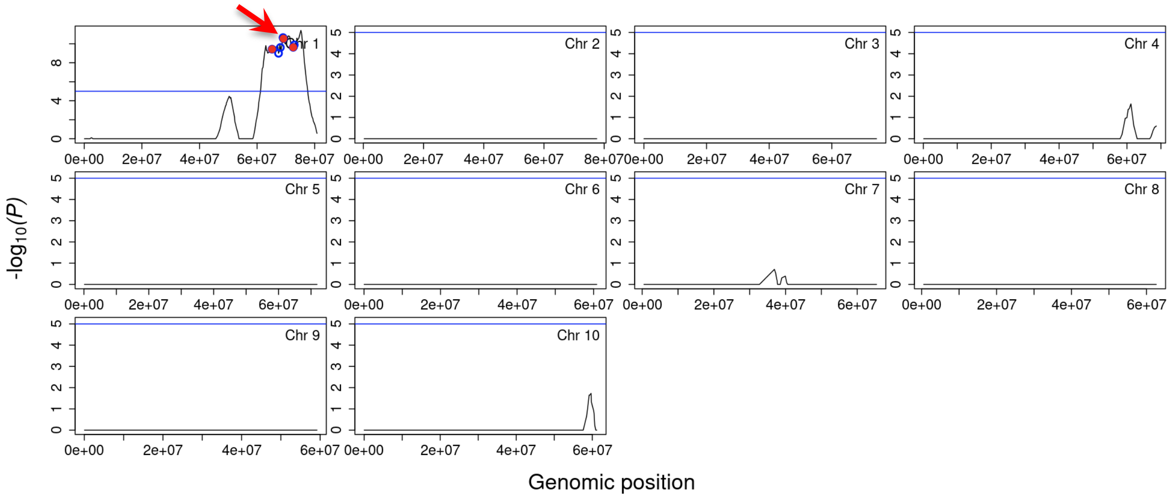


(E)

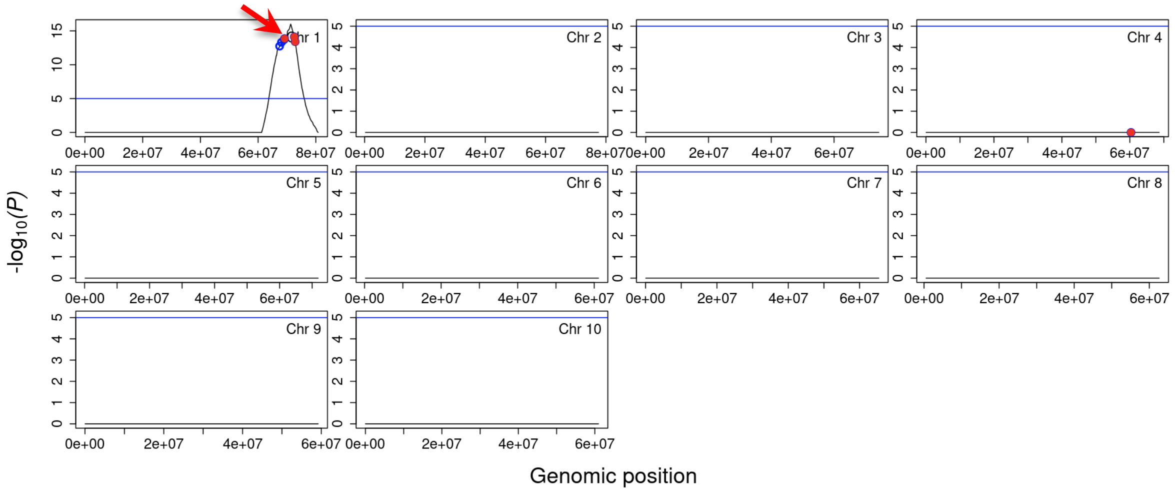


(F)

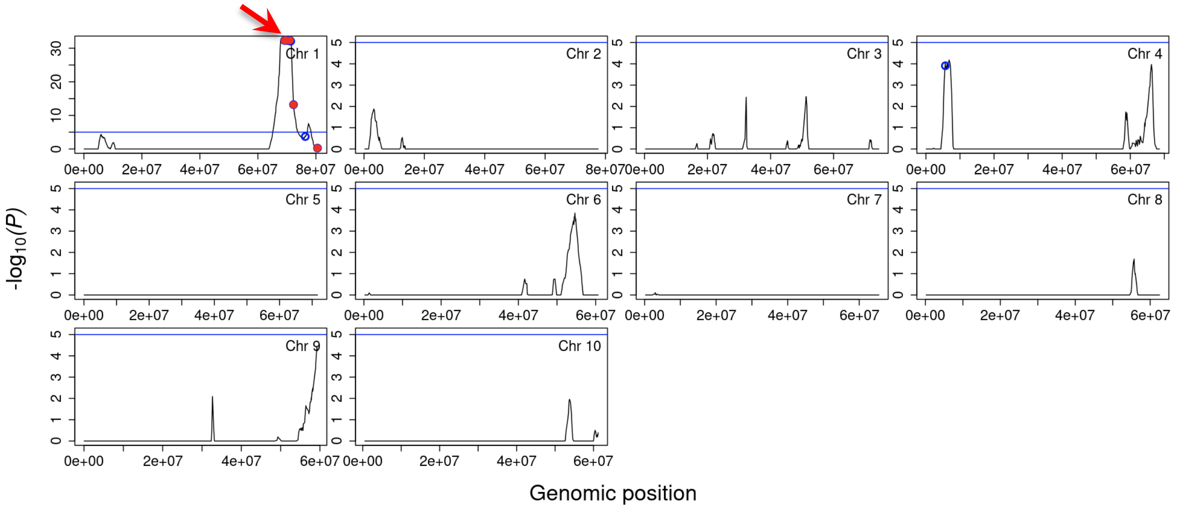


**Fig. S6 Segregation probability plots along the chromosomes for the *msd3* pools p6A_10Mb (A), p6B_5Mb (B), p14_2Mb (C), p21A_5Mb (D), p21B_10Mb (E), and p24_2Mb (F).**

Various window sizes were used to estimate the segregation regions for each F2 pool. The red arrow points to the causal mutation.
